# Deep-learning-assisted SICM for enhanced real-time imaging of nanoscale biological dynamics

**DOI:** 10.1101/2025.07.08.663539

**Authors:** Z. Ayar, M. Penedo, B. Drake, J. Shi, S. M. Leitao, I. Krawczuk, H. Miljkovic, A. Radenovic, J. Ban, V. Cevher, G. E. Fantner

## Abstract

Scanning Ion Conductance Microscopy (SICM) provides high-resolution, nanoscale imaging of living cells, but it is generally limited by a slow scan rate, making it challenging to capture dynamic processes in real time. To tackle this challenge, we propose an integrated data acquisition and computational framework that improves the temporal resolution of SICM by selectively skipping certain scan lines. A partial convolutional neural network (Partial-CNN) model is developed and trained on SICM images and their corresponding masks to reconstruct the complete images from the under-sampled data, ensuring the retention of structural integrity. This approach significantly reduces the image acquisition time (i.e., by 30-60%) without compromising quality, as validated through multiple quantitative metrics. Compared to conventional deep learning methods, the Partial-CNN demonstrates higher accuracy in reconstructing fine details and maintaining consistent height maps across skipped regions. We show that this method provides an increased temporal resolution and retains image fidelity, making it suitable for real-time dynamic SICM imaging and improving the smart scanning microscopy applications in time-resolved biological imaging.

**Graphical Abstract:** 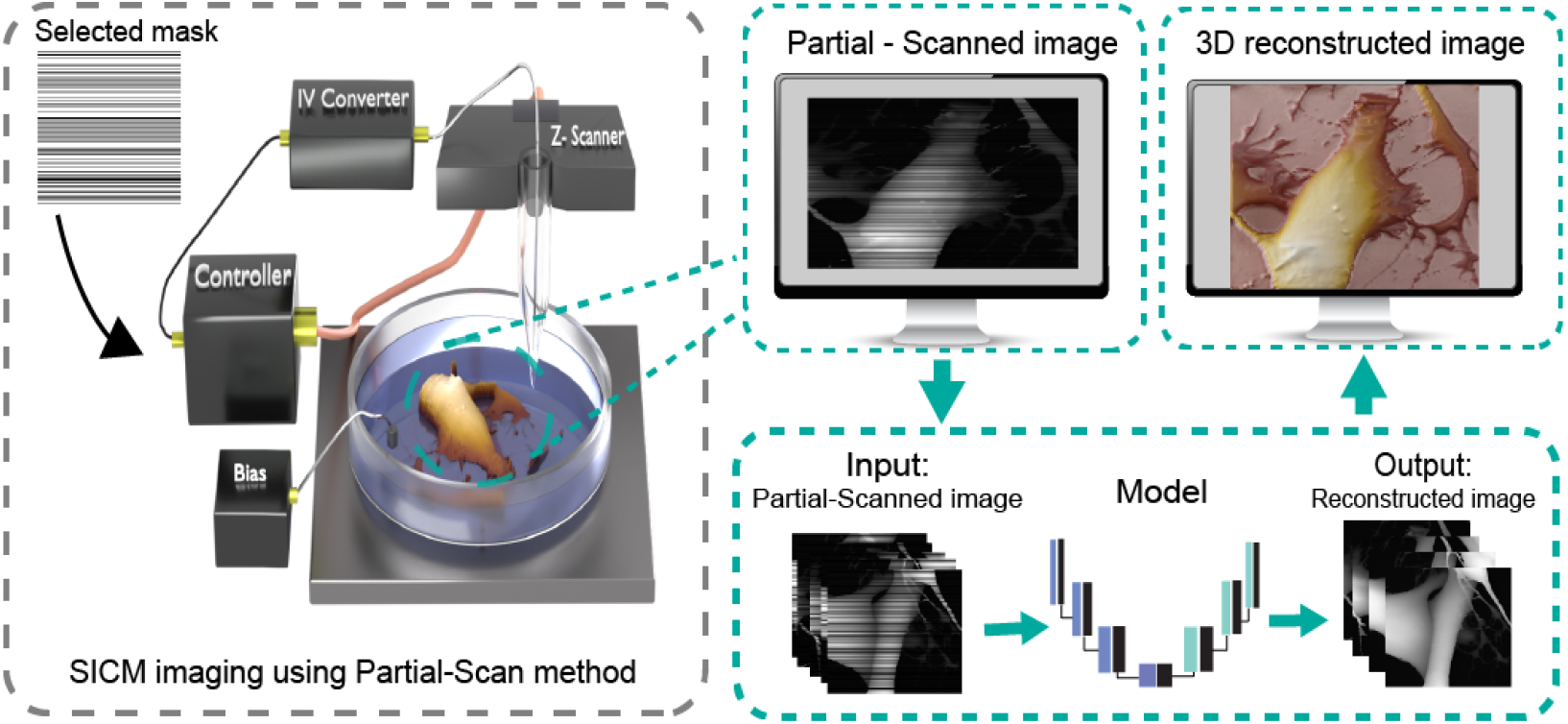

## 1. Introduction

Scanning ion conductance microscopy (SICM)^1^ is a subgroup of scanning probe microscopy (SPM) that is well-recognized for capturing high-resolution, non-invasive topographical images of live biological samples such as cells^1–4^. In SICM, the ionic current between a nanopipette and a sample is measured and used to track the sample surface, which allows for nanoscale cellular imaging without physically contacting the sample surface^1,5^. High lateral resolution (on the order of tens of nanometers) and axial resolution (< 5 nm)^5,6^, combined with the ability to image in a physiological environment (such as simple salt solution such as phosphate-buffered saline (PBS) to the more complex culture media such as DMEM, at 37°C with a CO_2_ supply), make this technique highly desirable for bio-imaging applications^7,8^.

The non-invasive nature of SICM offers distinct advantages over brightfield microscopy, not only in resolution but also in overcoming photosensitivity issues that can lead to phototoxicity in long-term live imaging^9^. Although atomic force microscopy (AFM)^10^ provides similar advantages for bio-imaging, it induces mechanical forces on the sample, potentially triggering mechanosensitive responses in cells. SICM has proven particularly useful in imaging fine structures in living cells, such as neurons^8^ and stem cells^11,12^, without any light-induced or mechanical stresses. Moreover, SICM shows extensive applications in detecting nano-mechanics^13,14^ and measuring stiffness^2,15^ of soft biological samples.

Despite its advantages, SICM is limited by the temporal resolution, especially when imaging the rapid movements of the cell membrane or the dynamics of fine membrane elements. Conventional SICM setups are slow, taking minutes to capture one frame. The scanning speed in SICM depends on several factors, including the mechanical design and resonant frequency of the piezoelectric actuators, the responsiveness of the feedback system, and the efficiency of current detection and processing circuits. High-speed SICM (HS-SICM) addresses these limitations and can reduce imaging times to a range of seconds per image by allowing an increased hopping rate from a few Hz (e.g., 20^16^ - 200^17^ Hz) up to several kHz^3,8,18^ on small scan sizes and flat samples. However, the required time and the distance that the probe has to move away from the surface before re-engaging in each hop, i.e., retract time and retract height, respectively, is considerably higher for some samples like neuron cells, which reduces the effective hopping rate.

To address SICM scanning speed limitation without requiring new equipment or hardware modifications, some previous studies focused on developing software-based solutions to reduce the scanning time. Takahashi et al. ^8^ developed an algorithm for the automation region of interest (AR mode) to limit the scanning area to the region with valuable data. Li et al. ^19^ used a pre-scanning low-resolution technique to detect the region with valuable data to decrease the scanning time. Gu et al. ^20^ used a target region focused (TRF) to skip the scanning of the background for an efficient scanning time. These methods are effective when there is a substantial area in the background to skip. For those images that contain layers of cells or stacks of biological samples, these methods do not reduce the scanning time. Additionally, edge detection techniques may face challenges in accurately predicting cell positions in the following frames during live imaging due to cell movement.

Emerging technologies like deep learning, specifically convolutional neural network (CNN)^21^, provide significant advancements in the performance of microscopy techniques, including segmentation^22^, super-resolution techniques^23^, and denoising^24^. Similarly, Liu et al.^23^ as well as Wu et al. ^25^ used a deep-learning approach respectively in AFM and SICM to enhance the resolution and generate high-resolution images from low-resolution inputs. These techniques allow scanning in a lower resolution (e.g., 32 × 32 pixels) and improve the image resolution (e.g., to 256 × 256 pixels) by an autoencoder, achieving results comparable to experimentally obtained high-resolution images. Although these models are effective, they miss fine features that are lost during the low-resolution scan, making the model unable to reconstruct the lost features.

This study introduces an alternative approach for data acquisition, which enhances imaging speed and reduces scanning time in SICM. The proposed technique selectively skips certain scan lines according to a predefined masking pattern, allowing a trained deep-learning model to reconstruct the missing data. We use two distinct models— convolutional neural networks (CNN)^21^ and partial convolutional neural networks (Partial-CNN)^26^— to demonstrate their effectiveness in reconstructing and inpainting images with skipped lines. We evaluate the performance of these models in reconstructing masked SICM images through visual inspection and quantitative metrics while comparing them with traditional linear interpolation methods. With this work, we contribute to developing smart microscopy techniques for scanning probe microscopy.

## 2. Results

### 2.1 Workflow of the Partial-Scan method

Figure 1 shows the overview of the workflow for the Partial-Scan method in SICM. A predefined mask determines which lines will be scanned and which will be skipped during the SICM partial scanning (Supplementary Data, Figure 1). The image and their corresponding masks are then fed to the model trained using a collection of previously recorded SICM images to reconstruct the skipped lines. Feeding the mask alongside the data improves accuracy by focusing the computation on the unmasked regions while also making the model lighter and more efficient for inpainting purposes of small datasets. The final reconstructed images are rendered to 3D RGB images to provide an intuitive visual representation.

**Figure 1:**
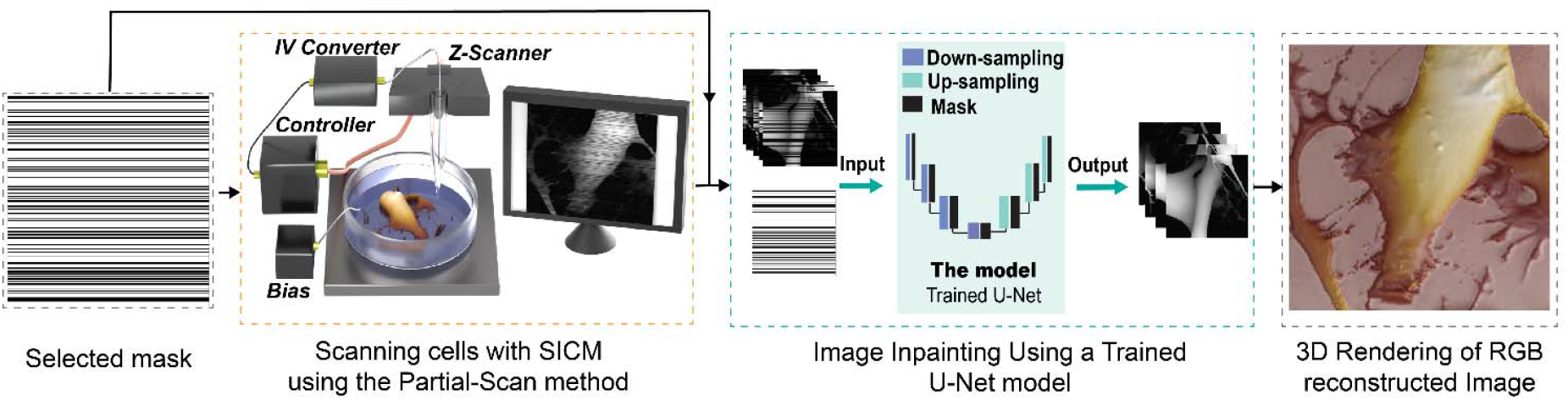
The workflow of the Partial-Scan method in SICM. After selecting the mask, the imaging was conducted according to the white lines (scanned lines). Both images and masks were fed to a pre-trained model to reconstruct the skipped lines. The image was visualized using a 3D RGB rendering to enhance the visual inspection.

### 2.2 Model performance

Two models were trained based on CNN and Partial-CNN using SICM images to compare their abilities to reconstruct skipped lines (Supplementary Figure S2). The Partial-CNN model has significantly fewer parameters to train due to the partial convolution in its layers, making it a less complex and more computationally efficient model. In contrast, the CNN model requires additional steps to reach optimal performance, while the Partial-CNN achieved an optimal reconstruction more quickly and is less prone to overfitting on a limited SICM data set (Supplementary Table S3, P < 0.001). We used a linear interpolation method to provide a minimum baseline reconstruction for model comparisons.

To evaluate the quality of the reconstructions, we first acquired fully scanned images and simulated the Partial-Scan method by removing the masked lines. This approach allowed us to compare the different reconstruction methods—linear interpolation, CNN, and Partial-CNN—against the original data, providing a clear benchmark for assessing their performance.

Figure 2.a illustrates the visual differences in reconstruction quality between linear interpolation, CNN, and Partial-CNN, highlighting the strengths and weaknesses of each approach in preserving the image details and structure. Figure 2.a.II demonstrates that linear interpolation cannot preserve the unscanned details. The reconstructed image using the CNN model (Figure 2.a.III) demonstrates overall high reconstruction quality; however, while it was able to capture most of the fine features, certain elements are lost during reconstruction in the area with file details (green arrows). The comparison between Figure 2.a.III and Figure 2.a.IV reveals that using Partial-CNN results in higher accuracy for capturing details and fine elements (green arrows). The observations (i.e., inside the enclosed blue dashed rectangle) show that the CNN model sometimes struggles to estimate the correct height map for the lost region and generates the wrong height value in the image.

**Figure 2:**
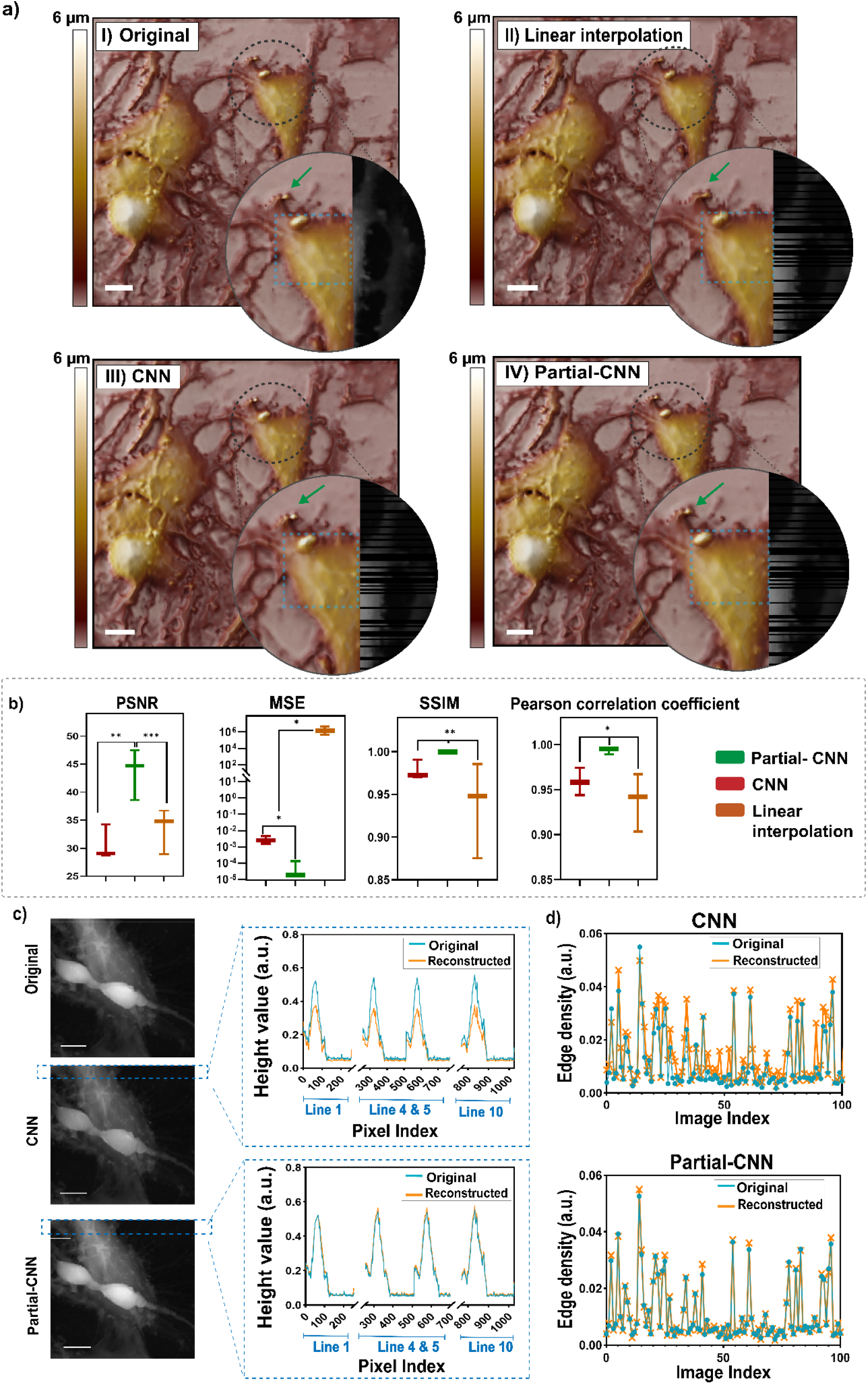
Comparison of reconstruction methods for SICM images (256 × 256 pixels) with 30% line masking; **a)** Visual comparison of (I)the original image and reconstructed image using (II) linear interpolation, (III) CNN, and (IV) Partial-CNN models. Linear interpolation lacks resolution, CNN misses capturing some features (green arrows) and creates incorrect height values (inside the blue rectangle). Scale bar = 10 µm. **b)** Statistical metrics (PSNR, MSE, SSIM, Pearson’s correlation coefficient) demonstrate a significant difference (p < 0.05) between CNN and Partial-CNN reconstructions, supporting previous observations. **c)** Visual inspection and height value analysis confirm that CNN struggles to maintain fidelity to the original image height values; however, Partial-CNN aligns closely with the original height values (scale bar = 10 µm); **d)** The canny method was used to calculate the edge density (number of pixels containing an edge normalized by the total number of pixels) for 100 images reconstructed using CNN and Partial-CNN. The result shows that CNN does not preserve the edges, and the method fails to keep the clarity of edges, while in Partial-CNN, it preserved all edges effectively.

The statistical analysis between CNN, Partial-CNN, and linear interpolation, shown in Figure 2.b, supports the visual observations, showing that the Partial-CNN performs better in image reconstruction compared to both other models. Metrics such as Peak Signal-to-Noise Ratio (PSNR), Mean Squared Error (MSE), Structural Similarity Index Measurement (SSIM), and Pearson’s correlation coefficient reveal statistically significant benefits of the Partial-CNN (p < 0.05). Although the CNN and linear interpolation performance in PSNR, SSIM, and Pearson’s correlation coefficient does not show significant performance, the MSE has a significant difference, showing that linear interpolation causes pixel-wise errors that leads to blurriness of reconstructed image.

The reconstructed images in Figure 2.c reveal that the CNN reconstruction cannot estimate the height values for the lost region compared to the Partial-CNN reconstruction. The height value when using Partial-CNN matches the original image, while the height value in the reconstructed lines is underestimated when using CNN. More investigation using the Canny method is performed to confirm the preservation of the edges. The calculation of the edge density of 100 images reconstructed by Partial-CNN and CNN is shown in Figure 2.d, which confirms lower preserved edges in images reconstructed using the CNN model compared to Partial-CNN.

### 2.3 Real-life image acquisition using the Partial-Scan method

The Partial-Scan method reduces the image acquisition time by reducing the number of scanned lines based on a predefined mask (e.g., 30% masking, Figure 3. a), where lines within the white regions of the mask are scanned, while black lines are skipped (Figure 3. b). To validate this approach, first, we image the fixed SH-SY5Y human neuroblastoma cells using the Partial-Scan method and then imaged the same region using a full-scan method. The comparison between the reconstructed images using the Partial-CNN model (Figure 3.c) and the original full-scan image (Figure 3.d) highlights the high efficacy and accuracy of the model in reconstructing the skipped lines (PSNR: 43, MSE: 1.5 ×10^−4^, SSIM: 0.99).

**Figure 3:**
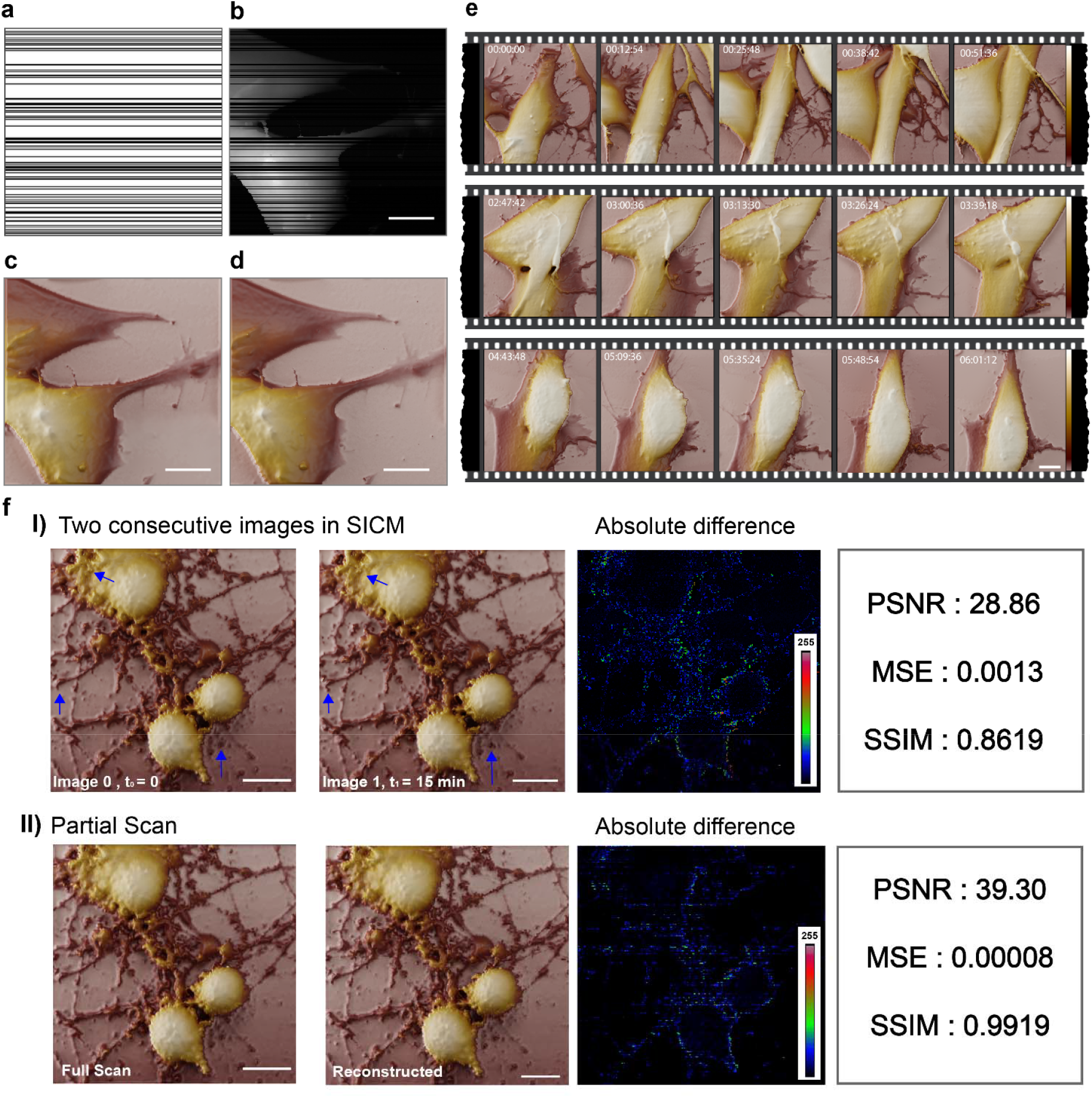
Evaluation of Partial-Scan method and image reconstruction in SICM on fixed and live cells using Partial-CNN; a) mask for skipped-line imaging (30% masking), b) partial scanned image, c) Reconstructed image using the Partial-CNN model, d) The full-scan image for reference, used to assess reconstruction accuracy, e) Time-lapse sections of live-imaging of neuroblastoma cells for more than 9 hours using skipped line method, f) Comparison of the absolute difference in I) two consecutive full-scan images in SICM, and II) in the Scan-Partial method shows that the uncertainty in the reconstruction based on Partial-CNN is lower than the inherent uncertainty of the instrument. Scale bars = 10 µm.

After validating the method on fixed samples, we applied it to image live SH-SY5Y neuroblastoma cells. The live cells were maintained in a custom-built incubator at 37°C with a controlled CO_2_ supply for over 9 hours. Figure 3.e presents sections from the time-lapse imaging of live cells, presenting the success of the Partial-CNN model in reconstructing skipped lines over extended imaging sessions. The Partial-Scan method reduces scanning time from 19 minutes 37 seconds to 13 minutes 48 seconds, achieving a 30% reduction. This time savings is critical for live-cell imaging, allowing for capturing more frames over time and enabling the observation of faster cellular movements and dynamics with greater accuracy (Supplementary Video S7).

It is shown in Figure 3.f to note that even in full-scan imaging of the same area, slight differences between two consecutive images can occur due to environmental factors, artifacts, or system noise. The comparison of images resulted from two consecutive full-scan and also between a full scan and Partial-scan method shows that metrics on reconstructed images (PSNR: 39.30, MSE: 8 × 10^−5^, SSIM: 0.9919) outperform the two consecutive full scan images (PSNR: 28.86, MSE: 0.0013, SSIM: 0.8619).

This indicates that the uncertainty of the model is lower than the inherent uncertainty of the SICM instrument.

### 2.4 Model’s break-point and masking effect

The performance evaluation of the Partial-CNN model across different masking percentages using PSNR, MSE, SSIM, and Pearson correlation coefficient in Figure 4.a reveals break points for the models calculated using segmented regression. Up to the primary break-point of 36% masking, the model performs well, maintaining high reconstruction quality with minimal artifacts and strong metrics across all evaluations. Between the primary break-point (36%) and the secondary break-point (55%), the model shows acceptable reconstruction with a gradual decline in all metrics, indicating challenges in accurately reconstructing fine details and maintaining structural integrity. Beyond 55% masking, the performance significantly deteriorated, with substantial loss of details and high-frequency information, making reconstruction unreliable (Supplementary Figure S4).

**Figure 4:**
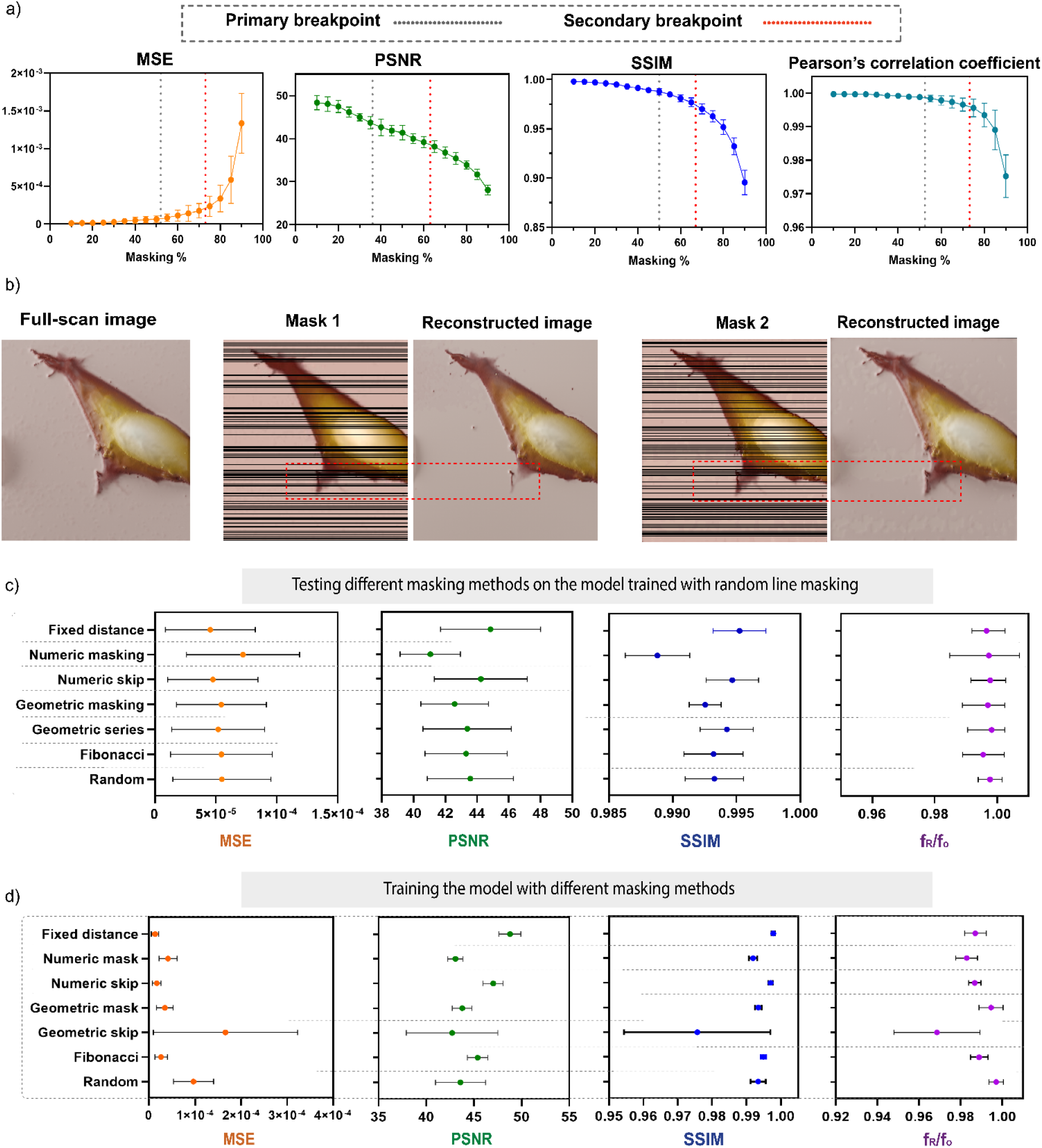
Model evaluations; a) Breaking point assessments show the primary break-point around 30% with the highest reconstruction potential according to all metrics. The metrics show acceptable reconstruction performance until 60%, with the possibility of some artifacts. b) Examples of the controlled-pattern masking methods generated using various logical patterns to assess the effect of masking on model performance, c) Evaluation of the potential of using different types of masks in the Partial-CNN model trained by random masking using the evaluation of PSNR, MSE, SSIM, and frequency content, the ratio of f_R_ - the frequency content of the reconstructed image-, and f_O_ - the frequency content of the original image. The result shows no significant difference. d) Performance comparison of the Partial-CNN models trained using different masking methods based on PSNR, MSE, SSIM, and frequency content. Although all the masking methods, except the Geometric mask, show no significant difference in PSNR, MSE, and SSIM, they all show lower frequency content compared to the random selection, which shows the effectiveness of the Random mask for training.

However, uncontrolled random masking could reduce the reconstruction quality if excessive information is lost in specific regions (Figure 4.b). To avoid data loss, we can limit the number of consecutive masked lines or train the model based on a controlled pattern mask, especially in higher masking percentages.

To assess if the masking pattern affects the image reconstruction quality, we generated several controlled masking patterns (Supplementary Figure S5), each generated based on a distinct logical approach (see Supplementary Table S6). Statistical analysis between the metrics in Figure 4.c reveals that using the different types of masking for a Partial-CNN model trained by Random mask shows no significant difference in the model performance, indicating the flexibility of this model for using any mask. We also evaluated whether the model trained using different controlled masking methods can provide an advantage over the Random line selection (Figure 4.d). Although the performance in PSNR, MSE, and SSIM shows no significant difference, except for the Geometric mask, the frequency content, which is the ratio of f_R_ (the frequency content of the reconstructed image), and f_O_ (the frequency content of the original image), reduces in all the other masking methods compared to Random mask. The results indicate that training the model with a specific pattern (e.g., Fixed distance) can lead to the loss of information in the frequency domain due to the fixed and repetitive pattern.

### 2.5 Using Partial-CNN to remove inherent scan artifacts

Beyond improving scanning speed and saving time in SICM, Partial-CNN models address additional challenges in SICM and, more broadly, scanning probe microscopies (SPM). Issues such as a contaminated medium or pipette overshooting from feedback delays can introduce artifacts, or “scars,” into the image, affecting its quality and usability. Traditional techniques like median line correction are often used to address these scars. Tools such as the open-source SPM image processing software Gwyddion use median-based methods to smooth out line defects by averaging neighboring lines, which can be effective for minor artifacts. However, as shown in Figure 5.a, median-based interpolation struggles to reconstruct these areas accurately when scarring is extensive. By considering the scarred regions as a mask (Figure 5.b), the Partial-CNN model effectively predicts and reconstructs the missing information (Figure 5.d) without introducing the blurriness often seen with median-based methods. The comparison between median line correction and Partial-CNN reconstruction highlights the Partial-CNN model’s advantage in preserving image clarity, showing its broader potential for image processing.

**Figure 5:**
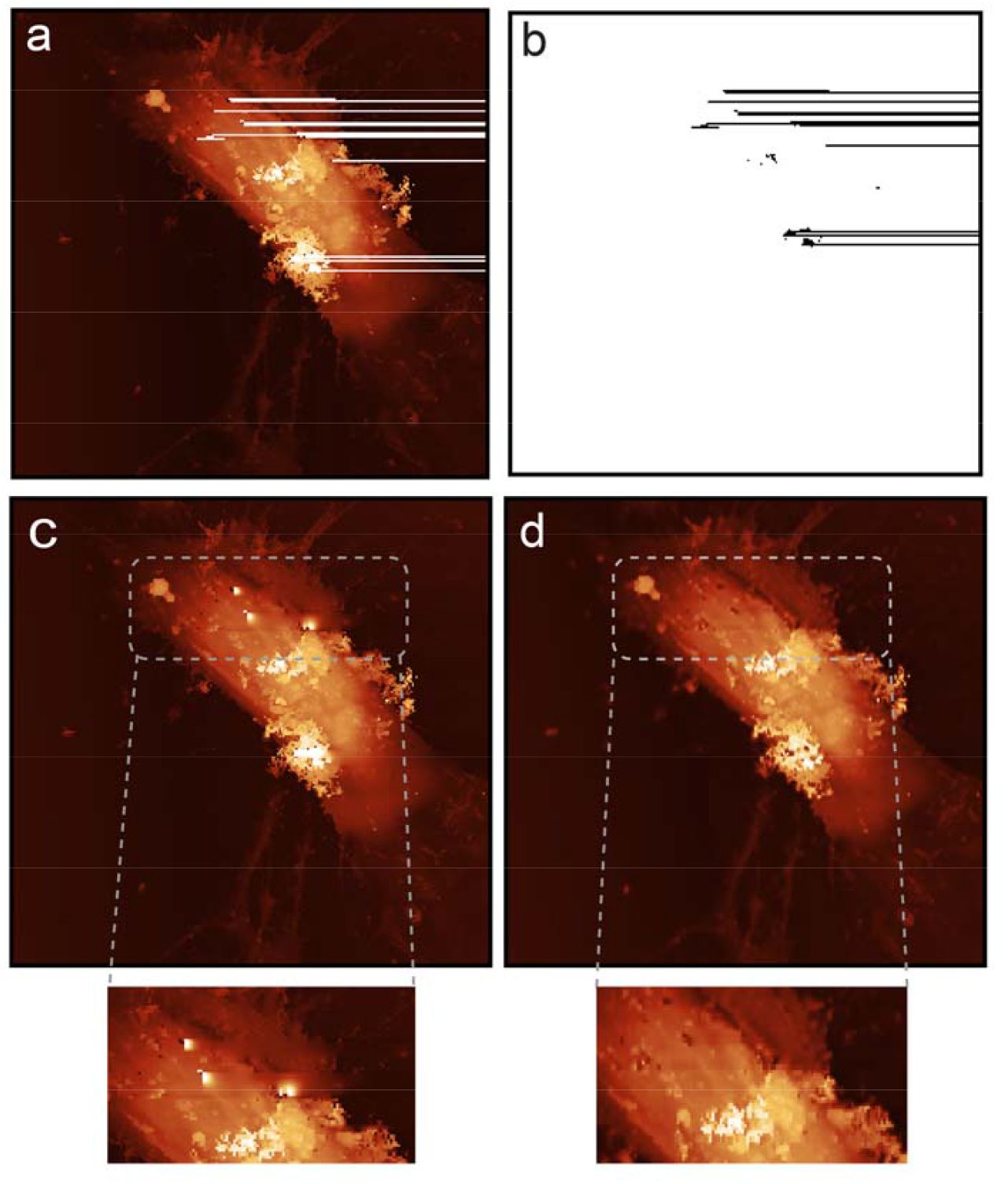
Extended application of the Partial-CNN model in scar correction; a) scarred image, b) extracted mask correlated with the scar pattern, c) reconstructed lines using the median of lines method (Gwyddion software), d) Reconstructed lines using Partial-CNN model.

## 3. Discussion

The scanning rate is a critical factor in scanning probe microscopy, especially in studies of dynamic phenomena in biological samples. In AFM, scanning rates have advanced rapidly due to hardware improvements and feedback control optimization, reaching an imaging rate of several Hz^27–30^. Alongside hardware developments, software-based solutions have explored subsampling methods for AFM to reduce imaging time^8,20,25^. Traditional compressed sensing methods^31–33^ have been employed successfully for subsampling and reconstructing AFM images, though they are limited by their reliance on mathematical models and a constrained number of parameters for optimization. However, deep learning allows us to train models with millions of parameters quickly, providing high performance in image reconstruction. However, deep learning reduces the control over model behavior due to the “black-box” nature of neural networks.

Our study demonstrates that the Partial-Scan method effectively reduces scanning time in SICM by selectively masking lines. The optimal masking level was found to be around 30%, offering a balance between increased temporal resolution and preserved image structure. Further tests confirmed that this masking level could be reliably increased to 60% if a small loss in image fidelity is acceptable. Training the model with random masks is recommended to retain flexibility when choosing varied line-skipping patterns.

Using a Partial-CNN results in an efficient inpainting and reconstruction of masked lines, maintaining high-frequency details and overall image quality. Metrics such as PSNR, MSE, SSIM, Pearson’s correlation coefficient, and height map analysis confirmed that Partial-CNN outperforms traditional CNN in this application. As shown in Figure 2, CNNs^34^ struggles with preserving the details, often introducing height value mismatches compared to the original images, while Partial-CNN^26^ preserves these fine details more effectively. This improvement is attributed to using the mask during training for Partial-CNN, which reduces the trainable parameters and focuses on the relevant image regions. The comparison of PSNR and other metrics in previous studies^31–33^ shows the significant difference between the traditional compress sensing methods and the current application of Partial-CNN in the reconstruction of the lines. Although the PSNR data of another study that uses a different neural network model (i.e., ISTA-Net) in reconstructing AFM images^35^ is reported close to our study, the Partial-CNN outperforms the mentioned model.

The evaluation between two images in a consecutive full scan of a fixed sample shows that there are minor differences between the two images in different consecutive scans due to the presence of artifacts in media, noise, and another environmental factors. The results in Figure 3.f show that the Partial-Scan is a robust and reliable method, showing lower absolute differences and higher performance (PSNR: 39.30, MSE: 8 × 10^−5^, SSIM: 0.9919) compared to the natural variations between two consecutive images in SICM (PSNR: 28.86, MSE: 0.0013, SSIM: 0.8619).

While this method demonstrates high reconstruction accuracy, certain limitations must be considered. Although pixel intensity comparisons verify the fidelity of partial-CNN reconstructions, minor height value misestimations may occur. We advise that images will be labeled as AI-enhanced reconstructions. It should also be closely monitored that skipping lines does not create a stepped pattern in the edges of the reconstructed images.

Future work can combine the Partial-Scan method with previous software solutions^8,20^ for background skipping^8^, which could further reduce scanning time by omitting background areas in each line and applying the Partial-Scan method only to regions of interest.

## 4. Conclusion

This study successfully improved the temporal resolution of SICM imaging by implementing a Partial-Scan method with a deep learning-based model trained to reconstruct the skipped lines. This technique applied advanced methods to develop Smart-SICM imaging as a subgroup of smart microscopy techniques in SPM. Utilizing a partial convolutional neural network (Partial-CNN) resulted in superior performance to traditional CNNs, achieving more accurate reconstructions with preserved detail. The time saved was directly related to the percentage of skipped lines, with 30% masking identified as an optimal balance, significantly enhancing temporal resolution while maintaining the structural fidelity of the sample, making it desirable for real-time live imaging of cells.

## 5. Methods

### Image acquisition using scanning ion conductance microscopy

A custom-built SICM setup was established and used for imaging^18^. The system consists of a custom Z-actuator driven by a home-built controller for feedback controlling fine movements and a stepper motor for the course movements. The Z-actuator covers a high range of scanning in Z with a maximum of 22 µm. An XY scanner (P-517 CL Piezo NanoPositioner, Physik Instrumente) with a 100 µm travel range was used for moving the sample in X and Y, which is driven by a low-voltage amplifier (E501 Piezo Controller System, Physik Instrumente). The XY scanner and Z actuator are assembled on an inverted microscope (Olympus IX71). A custom-made LabView software has been developed for instrument control. Nanopipettes were made using borosilicate (Sutter Instrument) with a CO2 laser puller (Model P-2000, Sutter Instruments) with an inner radius of 40 – 60 nm. To skip the lines, a mask will be selected. Then, the modified slow scan axis signal (Y-signal) is calculated based on the selected mask and sent to the SICM instrument to skip the determined lines. To avoid creep, the signal will be change through an increase in the Y-axis slope instead of using step-wise motion. Then, the sample is imaged with only the non-masked lines acquired during the scan, reducing the scanning time.

### Cell culture

SH-SY5Y human neuroblastoma cells were cultured in 3.5 cm Petri dishes (ibidi, 81156) using cell culture medium consisting of DMEM/F12 (Sigma-Aldrich, Gibco, 11320033) supplemented with 10% fetal bovine serum (FBS, Sigma-Aldrich, F7524), 1% penicillin-streptomycin (PenStrep, Gibco, 15140122), and maintained in a humidified custom-built incubator inside SICM with 5% CO_2_ at 37°C.

### Data preparation

A total of 815 single-channel, greyscale SICM images (256 × 256 pixels) were obtained from various cell types, including monkey kidney fibroblast-like cells (COS-7), human melanoma cells (SKMEL), mouse melanoma cells (B16−F1, ATCC), human cervical cancer cells (HeLa), human neuroblastoma cells (SH-SY5Y), and cortical neuron cells derived from opossum^36^. Ten percent of the images were set aside for testing, with the remaining split into training and validation sets (80% and 20%, respectively). Initial data observations ensured no leakage between sets. Predict data sets were used for real-life reconstruction. Due to the small dataset size, K-fold cross-validation^37^ was applied to reduce the risk of overfitting. This method divides data into three different sets (fold), with respective ratios for training, validation, and testing. Models were trained on each fold, and average metric values were reported.

### Masking methods

The images were masked using various techniques, with the masking percentage maintained between 30–35% of lines for model performance tests. Details of the controlled masking methods are provided in Supplementary Data, Table S6. In each method, white lines were scanned while black ones were skipped.

### Network Architecture and Implementation

Two U-Net-based^38^ models were developed to evaluate image inpainting, using consistent hyperparameters (batch size: 16, learning rate: 0.0005, alpha: 0.96) across both for optimization. The first model, CNN, employs an encoder-decoder structure with filters ranging from 64 to 512, incorporating 2D convolutional layers, max-pooling, and dropout (20%) for regularization (Supplementary Figure S2). The combined loss function is defined as:

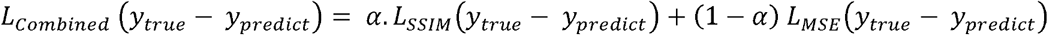

Where *L*_*Combined*_ is the total cost of the models, *y*_*true*_ is the original image, *y*_*predict*_ is the predicted image, is the learning rate, and *L*_*SSIM*_ is

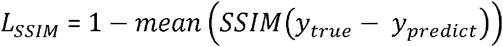

Which calculates the structural similarity between two images and, *L*_*MSE*_ is:

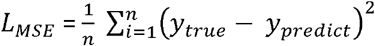

The second model, Partial-CNN ^26^, shares the CNN architecture but incorporates partial convolution layers and accepts both the masked image and mask as inputs. Element-wise multiplication before each convolution restricts processing to unmasked regions, with the mask dynamically updated during encoding and decoding. The loss function mirrors that of CNN. Models were trained on an NVIDIA A100-SXM4-80GB GPU. A linear interpolation technique scales images from 128×128 to 256×256 pixels as a baseline for reconstruction for visual inspections.

### Model Metrics

The model performance was evaluated using different mathematical metrics, including:

Mean Squared Error (MSE): the average of the squares of the differences between the original image and the reconstructed image.

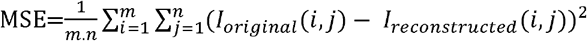

Where m and n are the dimensions of the image, and I is the pixel value in coordination with i and j. Peak Signal-to-Noise Ratio (PSNR): To evaluate image reconstruction quality.

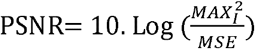

Where MAX_I_ is the maximum possible pixel value of the image, and MSE is the mean squared error between the original and reconstructed images.

Structural Similarity Index Measure (SSIM): To assess the perceptual similarity between two images based on the structural information, contrast, and luminance between the original and reconstructed images.

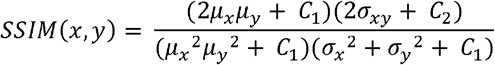

Where x is the original image, and y is the reconstructed image, μ_*x*_ and μ_*y*_ are the average (mean) value of x and y, σ_*x*_ and σ_*y*_ Are the variances of the original and reconstructed images, and and *C*_2_ are constants.

Edge density measurement: Canny edge detection method^39^ was used to detect the ratio of the edge pixels to the total pixel size in 100 images to determine the efficacy of the model in preserving the edges in the reconstructed image.

### Data Analysis

The preprocessing of images was handled using Gwyddion software. Skimage and Numpy Python libraries were used to calculate metrics (i.e., PSNR, MSE, SSIM, edge density). For statistical analysis, all data were analyzed using GraphPad Prism to perform one-way ANOVA, and t-test were used to compare the data groups. The 3D reconstruction of images was done using ImageJ and Blender 4.1 software. Model breakdown points is calculated based on segmented linear regression.

## Acknowledgments

This project has received funding from the European Union’s Horizon 2020 research and innovation programme under the Marie Skłodowska-Curie grant agreement No 945363. G.E.F., Z.A, M.P. sand B.F.D. also acknowledge, the Swiss Commission for Technology and Innovation under grant CTI-18330.1, the European Research Council under grant No. ERC-2017-CoG InCell, EPFL Center for Imaging under grant No. 563292, and ETH domain - ETH Open Research Data (ETH-ORD) under grant No. 563386. H.M. and A.R. acknowledges support from the European Research Council under grant no. 101020445—2D-LIQUID. Hereby, we acknowledge the Research Computing Platform (RCP)at EPFL for providing us with the computational facility.

## Notes

### Competing Interest Statement

The authors have declared no competing interest.

