## Supplementary document for "Deep-learning-assisted SICM for enhanced real-time imaging of nanoscale biological dynamics"

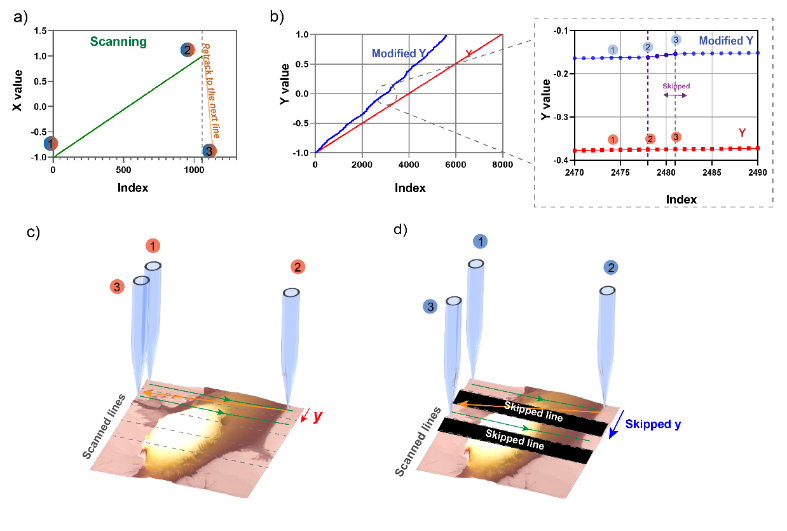

Supplementary Figure S1. The movement of x and y are precisely controlled by the XY scanner stage and its controller. Scanning in the x direction begins at point one (pixel 1) and proceeds to point 2 (the last pixel, e.g., 256). The stage then returns to the beginning of the next line at point 3. Typically, the y-axis moves at a constant linear speed (Figure 3.b) while the x-axis returns from point 2 to point 3. However, in the skip-line method, the y-axis movement follows the selected mask, resulting in a steeper y-axis slope. This allows for skipping more lines (e.g.,2y) within the same time frame that the x-axis returns to the next scanned line (from point 2 to point 3). This steeper movement reduces the overall scanning time without compromising the acquired resolution.

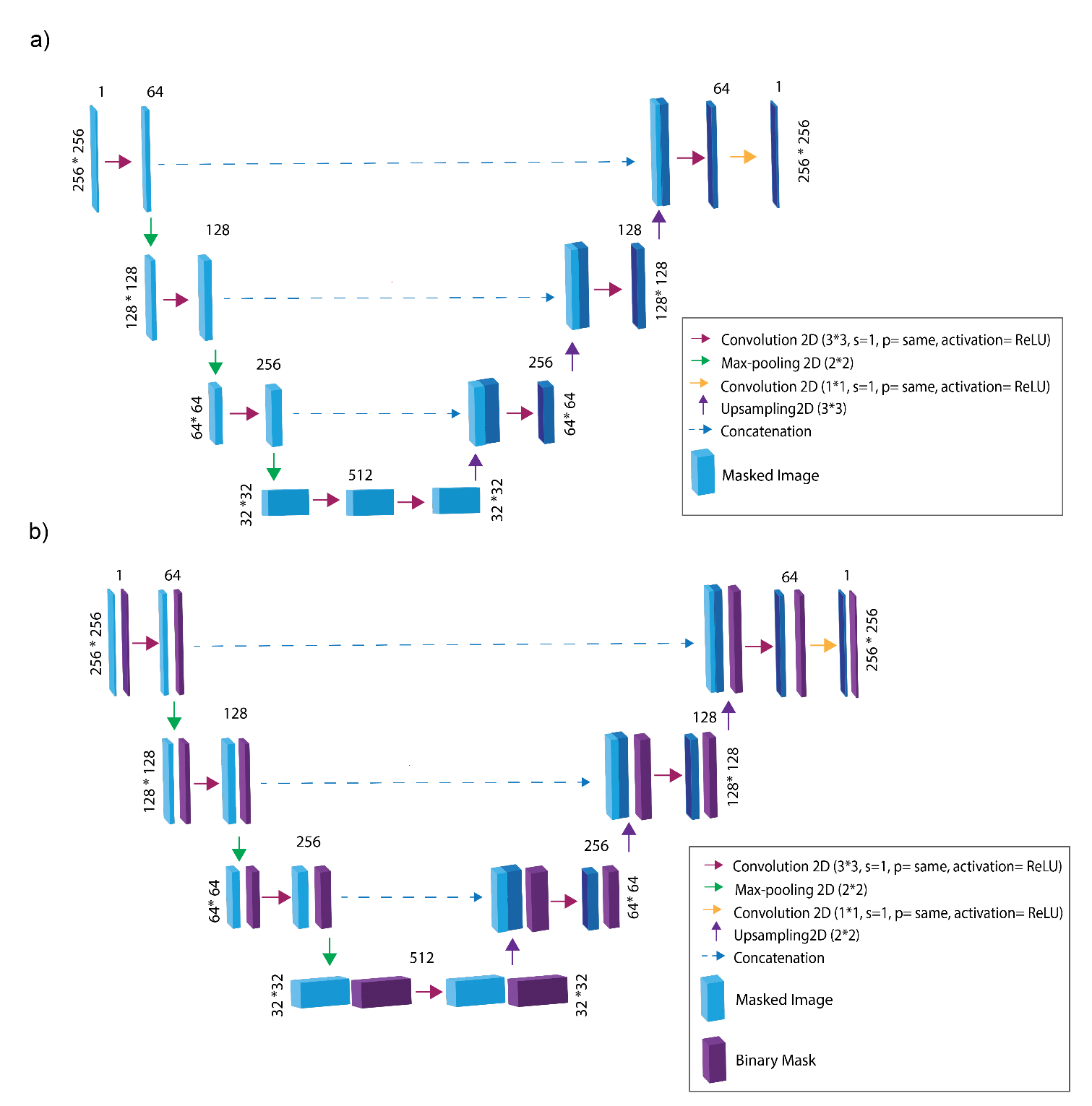

Supplementary Figure S2. The schematic of neural network models. a) Convolutional neural network (CNN), b) partial convolutional neural network (Partial-CNN)

Supplementary Table S3. Comparison of CNN and Partial-CNN model in consumption of resources.

| Metrics | CNN | Partial CNN |
| --- | --- | --- |
| Trainable parameters* | 16,453,697 | 2,403,448 |
| Training time | 7,505.33 ± 489.98 | 3,376.33 ± 131.18 |
| GPU Utilization (%) | 64.54 ± 1.54 | 72.25 ± 0.42 |
| GPU Memory Usage (GB) | 32.76± 0.76 | 32.10 ± 1.69 |
| Total Energy Consumption (J) * | 49,754.33 ± 3013.84 | 20,105.62 ± 474.22 |

*Significant difference (P < 0.001)

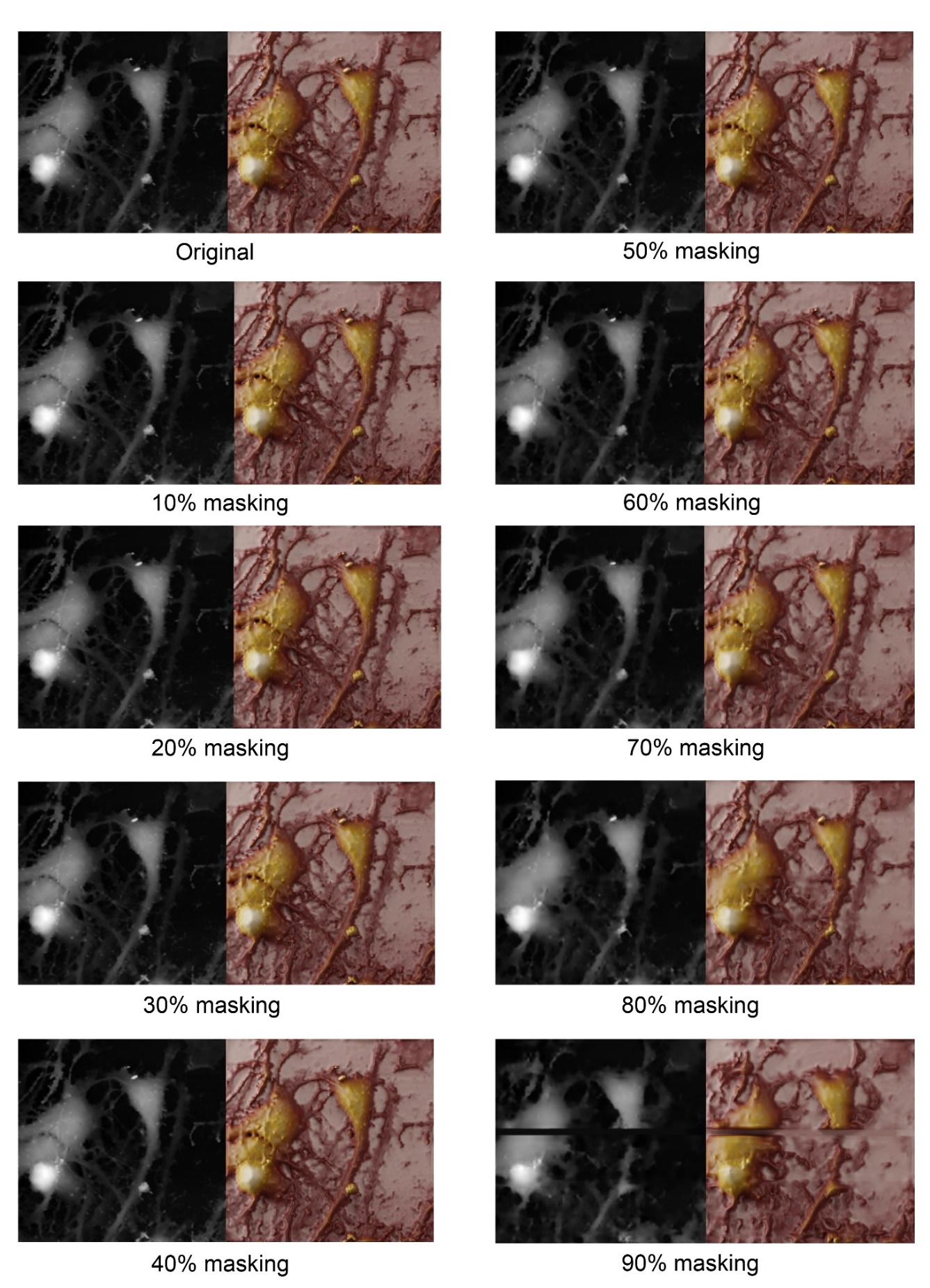

Supplementary Figure S4. The effect of masking percentage in reconstruction quality using Partial-CNN model shown in grayscale (model’s raw output) and 3D RGB rendered images.

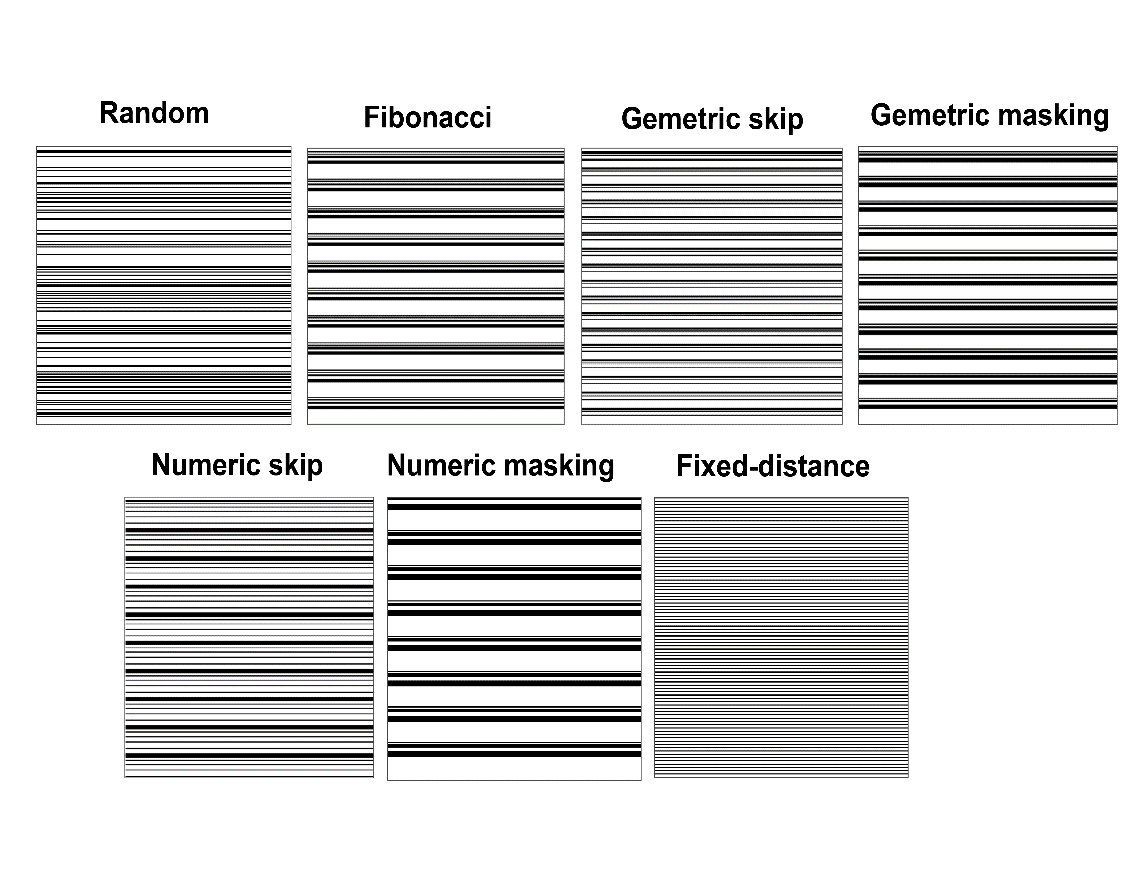

Supplementary Figure S5. Example of random masking with excessive consecutive masked lines, leading to significant information loss.

Supplementary Table S6. The controlled masking pattern strategies

| Masking Technique | Description | Series Formula | Pattern |
| --- | --- | --- | --- |
| Random | Random selection of horizontal lines with restriction of no masking more than 3 consecutive lines. |  |  |
| Fixed distance | Masking horizontal lines at regular intervals (Interval=3). | Interval = 3 | 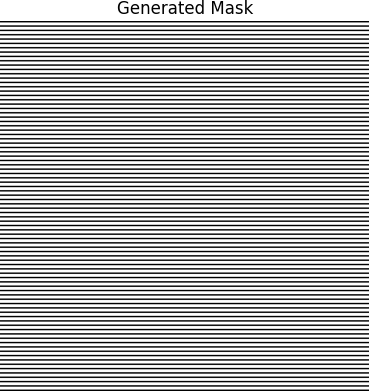 |
| Fibonacci | Masking lines based on Fibonacci sequence, alternating between masking and skipping Fibonacci number of lines. | $T_{n}= T_{n-1}- T_{n-2}$  Max. num = 3, repeat = 10 | 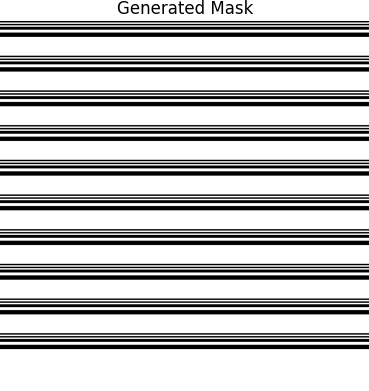 |
| Geometric mask | Masking lines using geometric series to determine skips after masking each line. | $T_{n}=a.r^{(n-1)}$  a=1, r=2, Max. n = 3, repeat = 30 | 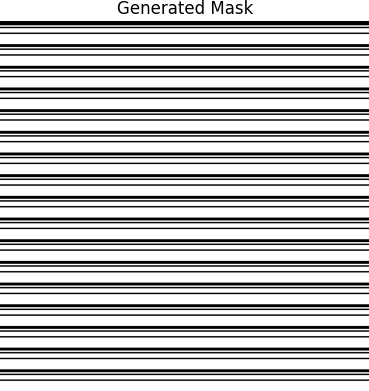 |
| Geometric skip | Masks and skips lines using a geometric progression with increasing gaps and masked lines. | $T_{n}=a.r^{(n-1)}$  a=1, r=2, Max. n = 3, repeat = 11 | 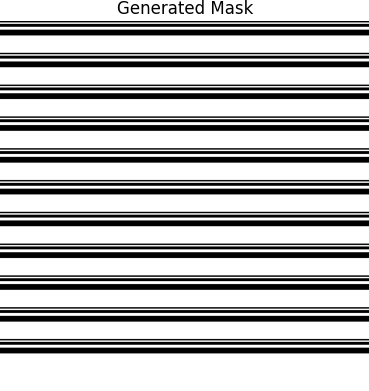 |
| Numeric mask | Masking lines using an arithmetic numeric series to determine skips after masking each line. | $T_{n}= a+\left( n-1 \right).d$  a=1, d=2, Max. num = 3, repeat = 10 | 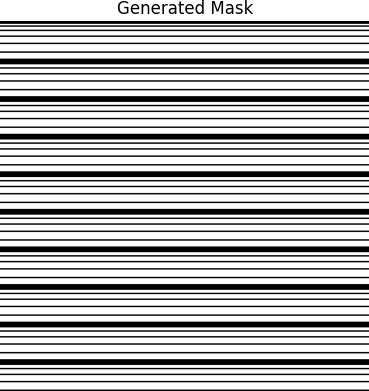 |
| Numeric skip | Masks and skips lines using a numeric progression, with increasing gaps and masked lines. | $T_{n}= a+\left( n-1 \right).d$  a=1, d=2, Max. num = 3, repeat = 8 | 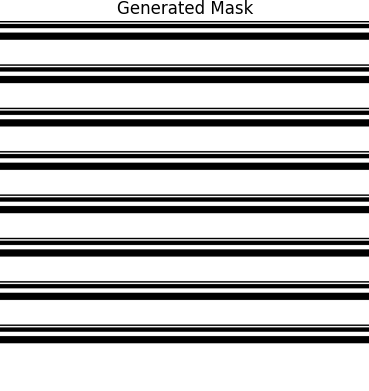 |

Supplementary Video S7. The live time-lapse for more than 9 hours of neuroblastoma cells
